# Assessment of Water Stress Tolerance in Mungbean Induced by Polyethyline Glycol

**DOI:** 10.1101/872663

**Authors:** Mohammad Rafiqul Islam, Mohammad Sohidul Islam, Bikash Chandra Sarker, Ashraful Alam, Masuma Akhter, Mohammad Jahangir Alam, Hirofumi Saneoka, Murat Erman, Ayman EL Sabagh

## Abstract

Water scarcity is a common hindrance of crop production in the globe especially in dry regions due to low precipitation along with high air temperature. Mungbean is a valuable pulse crop faced considerable amount of water shortage during its growing periods (March-May). Thus, the study was carried out to find out the variability and diversity present in the mungbean genotypes based on genetic variation of germination and seedling growth traits induced by polyethyline glycol (PEG-6000) stress (−0.7, −1, −2 and −4 bar). Seeds of each genotype was placed in a 9 cm diameter petridish containing sand bed that moistened by respective prepared PEG solutions and allowed to grow up to 10 days after placement. The result showed that the phenotypic coefficient of variation was significantly greater than the genotypic coefficient of variation for the entire studies traits, representing that these traits was influence by the environmental factors. The heritability showed moderate to high (24.26%-99.19%) under the observed traits indicating that these traits were under the control of additive genetic effects. The clustering pattern exhibited that cluster III were the largest ones with eleven genotypes. In case of contribution of different traits, the first five components were found to contribute for 98.19 % of the total variation while only first two components contributed for 82.12 % variation. The greater inter-cluster distance was found among the cluster I with II, III, IV and V. The maximum intra-cluster distance was observed in the cluster II (0.12). The genotypes in the cluster V recorded for greater performance by giving the highest mean value for germination, coefficient of germination, speed of germination, vigor index, shoot length and root length. This mean that genotype under cluster V have any gene or mechanism which responsible for tolerant higher level of water stress. The results of the study can be caring for establishing assortment criteria for water stress tolerant mungbean genotypes under breeding program.

## Introduction

With the progress of global climate change, crops are facing more challenges due to different biotic and abiotic stress. Different abiotic stresses viz., drought, salinity, heat, cold, water logging etc. had an adverse effect on crop growth and developments and ultimately on the final crop production. Among them, drought stress is one of the burning issues that limiting crop performances. Scarcity of water limits plant survival and early seedling growth by delaying its establishment or decreasing the final germinability [1]. Among different growth processes, germination of seeds and seedling growth at early stages are considered as the most critical phases for seed establishment, determining successful crop production [2].

Mungbean (*Vigna radiata* L. Wilczek) is an important short duration pulse crop that grows worldwide is an important source of plant protein. Its absolute dependence on monsoon rains for moisture in combination with rapidly withdrawing rainfall is a barrier for normal physiological processes of growth and development [3]. Incidence of water stress in different scale during the crop growth period affect the number of a physiological and metabolic process like germination, growth, photosynthesis, respiration, nutrient metabolism, etc, as a result, leading to poor growth and yield [4]. Thus there is a great require for drought-resistant varieties that could survive limited soil moisture stress and produce a better yield. Variability present in the diverse germplasm pool inadequately utilized in different studies. Consequently, genotypes performance under drought stresses poorly studied.

Drought tolerance has a complex character that difficult to solve and involves different approaches and methods. Creating new races or genotypes could be facilitated by selection with higher resistance. The assortment of parents along with superior drought tolerance is critical in dry environments [5, 6]. Therefore, there have been attempts to determine the degree of tolerance with a single parameter have limited value because of the diversity of the factors and their relations contributing to drought tolerance under field conditions [7]. Studies on water potential enabled the recognition of genotypes appropriate for growing under water stress situations. Genotypes that are found to germinate under reduced water potential do not usually fail to germinate and establish into seedlings. However, changes in water potential, through the use of high molecular weight substances, like polyethylene glycol (PEG) cause osmotic shock which is one of the components of the drought, added to the medium for seed germination can enable the identification of genotypes suitable for growing under water stress. Therefore, the present study has been carried out to estimate the variability and diversity present in the mungbean germplasm.

## Materials and methods

Thirty-three mungbean genotypes including most of the popular varieties and advanced lines were collected from Bangladesh Agricultural Research Institute (BARI), Bangladesh Institute of Nuclear Agriculture (BINA) and Bangabandhu Sheikh Mujibur Rahman Agricultural University (BSMRAU), Bangladesh. The list of genotypes is presented in Table 1.

**Table 1.** The name of the mungbean genotypes with their pedigree used in the present study

| SL No | Genotypes | Pedigree/Parents | Remark | Year released/ Develop |
| --- | --- | --- | --- | --- |
| 1. | BARI Mung-1 (Mubarik) | Advance line of Mung 7706 (India) | V | 1982 |
| 2. | BARI Mung-2 (Kanti) | VC 2768A × VC 6141-36 (7715) | V | 1987 |
| 3. | BARI Mung-3 | Sonamung(Local) × BARI Mung- 2 (BMX-842243) | V | 1996 |
| 4. | BARI Mung-4 | Sonamung(Local) × BARI Mung- 2 (BMX-841121) | V | 1996 |
| 5. | BARI Mung-5 | VC- 2768B × VC-6141-36 (NM-92) (AVRDC) | V | 1997 |
| 6. | BARI Mung-6 | NM-36 × VC- 2768A (NM-94) (AVRDC) | V | 2003 |
| 7. | BARI Mung-7 | VC-3960A-88 × VC-6173C (BMX-97024-13) | V | 2015 |
| 8. | BARI Mung-8 | Local Selection (LM-101) | V | 2015 |
| 9. | BINA Mung-6 | VC-6173-10 collected from AVRDC | V | 2005 |
| 10. | BINA Mung-7 | BINA Mung-2 applying EMS mutagen | V | 2005 |
| 11. | BINA Mung-8 | MB-149 (Irradiated with 400 Gy dose of gamma ray) | V | 2010 |
| 12. | BU Mung-2 | VC 2768A x VC 6141-36 {AVRDC ID- VC-6370 (30-65)} | V | 2001 |
| 13. | BU Mung-4 | GK7 (AVRDC, Taiwan) | V | 2006 |
| 14. | BMX-01019-1 | 128001 × NM-94 | AL | 2001 |
| 15. | BMX-01007 | BARI Mung-3 × BARI Mung-5 | AL | 2001 |
| 16. | BMX-01015 | NM-94 × BARI Mung-3 | AL | 2001 |
| 17. | BMX-030011-6 | BMX-97004-1 × BARI Mung- 5 | AL | 2003 |
| 18. | BMX-04005-3 | BMX-97004-1 × BMX-97014-3 | AL | 2004 |
| 19. | BMX-05001 | BARI Mung- 5 × BARI Mung- 6 | AL | 2005 |
| 20. | BMX-05004 | BARI Mung- 6 × BARI Mung- 5 | AL | 2005 |
| 21. | BMX-05009 | BMX-94002-2 × BARI Mung- 6 | AL | 2005 |
| 22. | BMX-050012-6 | BMX-99002-2 × BARI Mung- 5 | AL | 2005 |
| 23. | BMX-06001 | BARI Mung- 5 × VC-6469-12-1-4A | AL | 2006 |
| 24. | BMX-06007 | BARI Mung- 5 × BMX-97024-8 | AL | 2006 |
| 25. | BMX-07007-4 | BMX-00014 × Nilphamari local | AL | 2007 |
| 26. | BMX-07009-10 | BARI Mung- 6 × BMX-00016 | AL | 2007 |
| 27. | BMX-08009-7 | BARI Mung- 6 × BAU Mung-2 | AL | 2008 |
| 28. | BMX-08010-2 | BARI Mung-6 × BMX-9902-2 | AL | 2008 |
| 29. | BMX-08011-8 | BARI Mung- 2 × BMX-9902-2 | AL | 2008 |
| 30. | BMX-08011-2 | BARI Mung- 2 × BMX-9902-2 | AL | 2008 |
| 31. | BMX-09009-6 | Nilphamari local × BMX-9902-2 | AL | 2009 |
| 32. | BMX-97024-13 | VC-3960A-88 × VC-6173C | AL | 1997 |
| 33. | VC-2764B | Advance line of Mung(AVRDC) | AL | - |

V= Variety, AL= Advance line

These genotypes were evaluated for drought tolerance imposed by five different levels of water potential (0, −0.7, −1, −2 and −4 bar) induced by Polyethylene Glycol (PEG). The trial was conducted at the Laboratory of Agronomy, Hajee Mohammad Danesh Science and Technology University (HSTU), Dinajpur, Bangladesh during 2016. The trial was set up in two factors completely randomized design (CRD) with three replications. Stress levels were considered as Factor A while genotypes were regarded as Factor B. Twenty (20) seeds of each genotype were planted on a 9 cm diameter Petridish containing sand bed that moistened by respective prepared PEG solutions of −0.7, −1, −2 and −4 bar. Thereafter, 5 ml of the solution of respective treatment were applied every day in each genotype of the particular replications. Seedlings were allowed to grow up to 10 days after the placement of germination. The solution of different concentrations of Polyethylene Glycol (PEG 6000) was calculated as described by Michel and Kaufmann [8] which equation as follows-Water potential (bar index) = (−1.18× 10^−2^) ×C - (1.18× 10^−4^)×C + (2.67 × 10^−4^) ×C×T + (8.39 × 10^−7^) × C^2^T Where C=PEG concentration, T=Temperature (centigrade).

After the end of 10th day, observations were recorded on the germination related traits such as-Germination percentage (GP), Speed of germination (SG), Co-efficient of germination (CEG), Vigor Index (VI) and Mean germination time (MGT) and seedling traits such as shoot length (SL), root length (RL), shoot fresh weight (SFW), shoot dry weight (SDW), root fresh weight (RFW) and root dry weight (RDW). GP was calculated by using the following formulae [9] and measured according to the International Seed Testing Association (ISTA), [10]. SG was calculated according to Maguire’s equation [11]. CEG was calculated as the formulae given by Copeland [12]. VI was calculated by using the formula of Baki and Anderson [13]. MGT was calculated by the following equation proposed by Moradi et al., 2008 [14]. The data for the final germination percentage was analyzed, after arcsine transformation [15].

### Statistical analysis

For statistical analysis, the values of germinating percentage were transformed to arcsine while values of other traits done by log transformation due to data contain zero value at higher stress levels. The data were analyzed by partitioning the total variance with the help of a computer using R software program [16] (R Core Team, 2019) following the basic procedure outlined by Gomez and Gomez [17]. The means separations were done by LSD at 0.05 levels of probability to determine the statistical differences between means when the F value was significant. MS Excel software program was used for drawing the diagrams. Multivariate analysis was done for grouping the mungbean genotypes based on stress tolerance. The data were also subjected to analysis according to Mahalanobis’ D2-statistics. The intra and inter cluster distance, cluster mean and involvement of each character to the divergence were measured as described by Singh and Chaudhary [18].

## Results and discussion

### Variability estimates of different germination and seedling related traits of mungbean genotypes

Variability in the respective genotypes is the pre-requisite for undertaking a varietal development program [19]. Therefore, analyzing the character and extent of the heritable genetic variation existent in the genotypes is essential. The magnitude of the genotypic and phenotypic coefficient of variation, heritability (h2b) with their genetic advance and genetic gain for germination and seedling characters of 33 mungbean genotypes have been presented in Table 2.

**Table 2.**
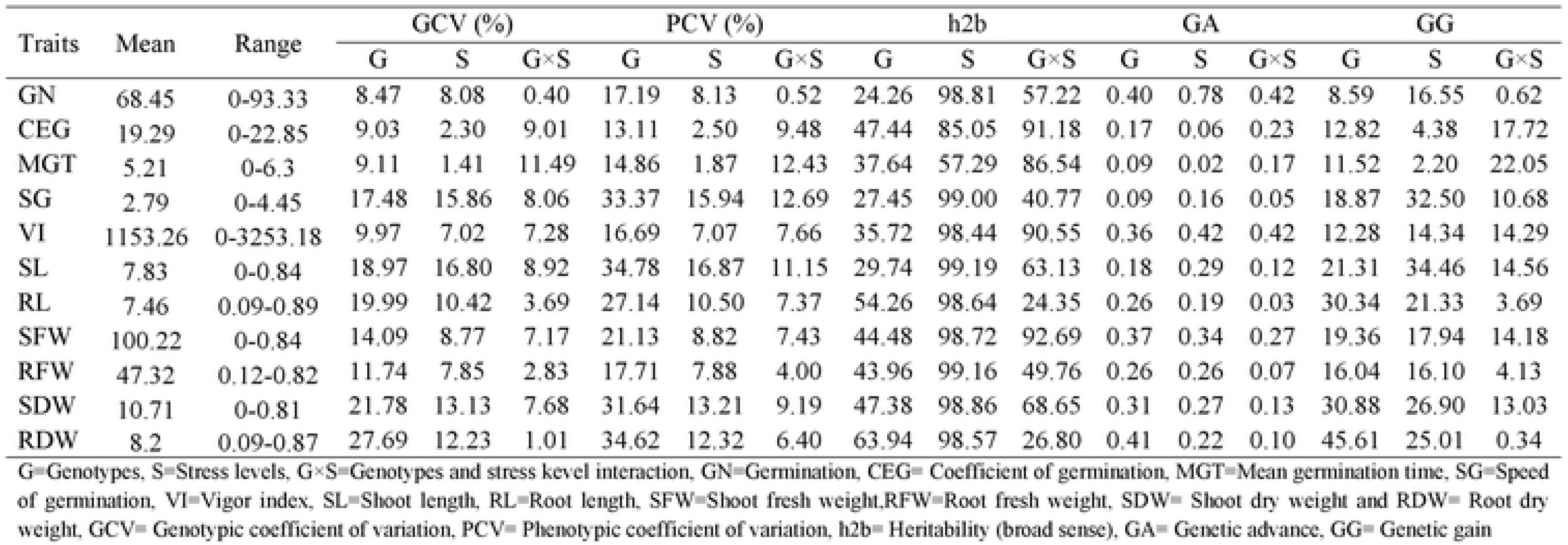
Variability estimates of different germination and seedling related traits

| Traits | Mean | Range | GCV (%) |  |  | PCV (%) |  |  | h<sup>2</sup>b |  |  | GA |  |  | GG |  |  |
| --- | --- | --- | --- | --- | --- | --- | --- | --- | --- | --- | --- | --- | --- | --- | --- | --- | --- |
|  |  |  | G | S | G×S | G | S | G×S | G | S | G×S | G | S | G×S | G | S | G×S |
| GN | 68.45 | 0-93.33 | 8.47 | 8.08 | 0.40 | 17.19 | 8.13 | 0.52 | 24.26 | 98.81 | 57.22 | 0.40 | 0.78 | 0.42 | 8.59 | 16.55 | 0.62 |
| CEG | 19.29 | 0-22.85 | 9.03 | 2.30 | 9.01 | 13.11 | 2.50 | 9.48 | 47.44 | 85.05 | 91.18 | 0.17 | 0.06 | 0.23 | 12.82 | 4.38 | 17.72 |
| MGT | 5.21 | 0-6.3 | 9.11 | 1.41 | 11.49 | 14.86 | 1.87 | 12.43 | 37.64 | 57.29 | 86.54 | 0.09 | 0.02 | 0.17 | 11.52 | 2.20 | 22.05 |
| SG | 2.79 | 0-4.45 | 17.48 | 15.86 | 8.06 | 33.37 | 15.94 | 12.69 | 27.45 | 99.00 | 40.77 | 0.09 | 0.16 | 0.05 | 18.87 | 32.50 | 10.68 |
| VI | 1153.26 | 0-3253.18 | 9.97 | 7.02 | 7.28 | 16.69 | 7.07 | 7.66 | 35.72 | 98.44 | 90.55 | 0.36 | 0.42 | 0.42 | 12.28 | 14.34 | 14.29 |
| SL | 7.83 | 0-0.84 | 18.97 | 16.80 | 8.92 | 34.78 | 16.87 | 11.15 | 29.74 | 99.19 | 63.13 | 0.18 | 0.29 | 0.12 | 21.31 | 34.46 | 14.56 |
| RL | 7.46 | 0.09-0.89 | 19.99 | 10.42 | 3.69 | 27.14 | 10.50 | 7.37 | 54.26 | 98.64 | 24.35 | 0.26 | 0.19 | 0.03 | 30.34 | 21.33 | 3.69 |
| SFW | 100.22 | 0-0.84 | 14.09 | 8.77 | 7.17 | 21.13 | 8.82 | 7.43 | 44.48 | 98.72 | 92.69 | 0.37 | 0.34 | 0.27 | 19.36 | 17.94 | 14.18 |
| RFW | 47.32 | 0.12-0.82 | 11.74 | 7.85 | 2.83 | 17.71 | 7.88 | 4.00 | 43.96 | 99.16 | 49.76 | 0.26 | 0.26 | 0.07 | 16.04 | 16.10 | 4.13 |
| SDW | 10.71 | 0-0.81 | 21.78 | 13.13 | 7.68 | 31.64 | 13.21 | 9.19 | 47.38 | 98.86 | 68.65 | 0.31 | 0.27 | 0.13 | 30.88 | 26.90 | 13.03 |
| RDW | 8.2 | 0.09-0.87 | 27.69 | 12.23 | 1.01 | 34.62 | 12.32 | 6.40 | 63.94 | 98.57 | 26.80 | 0.41 | 0.22 | 0.10 | 45.61 | 25.01 | 0.34 |

G=Genotypes, S=Stress levels, G×S=Genotypes and stress level interaction, GN=Germination, CEG= Coefficient of germination, MGT=Mean germination time, SG=Speed of germination, VI=Vigor index, SL=Shoot length, RL=Root length, SFW=Shoot fresh weight, RFW=Root fresh weight, SDW= Shoot dry weight and RDW= Root dry weight, GCV= Genotypic coefficient of variation, PCV= Phenotypic coefficient of variation, h<sup>2</sup>b= Heritability (broad sense), GA= Genetic advance, GG= Genetic gain

### Genotypic and Phenotypic Coefficient of Variation

The genotypic coefficient of variation (GCV) and phenotypic coefficient of variation (PCV showed wide variation for most of the characters under the study. The difference between PCV and GCV values reflects the influence of environmental effect on the character, in which the influence of the environment should be taken care of for a precise selection of material for the improvement of the crop [20]. Regarding the results, the GCV ranged from 8.47 to 27.69; 1.41 to 16.80 and 0.40 to 11.49, respectively in genotypes, stress levels, and their interaction. Similarly, the PCV ranged from 13.11 to 34.78; 1.87 to 16.87 and 0.52 to 12.69, respectively in the same treatments. However, the GCV was highest for RDW (27.69) followed by SDW (21.78) while the traits like GN (8.47) gave the lowest in case of genotypes. In stress level, SL gave the higher GCV (16.80) followed by SG (15.56) whereas the minimum was in MGT (1.41). In genotypes and stress kevel interaction, MGT exhibited the highest GCV value (11.49) and least was in GN (0.40). Considering the genotypes, the PCV was highest for SL (34.78) followed by RDW (34.62) and SDW (31.64) while the lowest value was in CEG (13.11). The traits like SL exhibited the highest PCV (16.78) followed by SG (15.94) in respect of stress level and the minimum was in MGT (1.87). In view of genotypes and stress kevel interaction, SG exhibited the highest PCV value (12.69) followed by MGT (12.43) and SL (11.15) whereas the last values of PCV were obtained for traits like GN (0.52. The estimated data revealed that the PCV was higher than the GCV for all the studied traits which indicate that there was some environmental influence on these traits. These results were also line with the findings of Khajudparn and Tantasawat, 2011; Garg et al., 2017; Garje et al., 2013; Gadakh et al,, 2013; Kumar et al., 2013; Degefa et al., 2014 [21–26]. The lowest values of PCV and GCV in respective traits followed by the higher values, indicating these to be the lowest variable characters [27].

### Heritability

Heritability refers to the ratio of phenotypic variance and genotype variance, i.e., heritable [20]. The results of heritability (broad sense) estimates have been presented in the Table 2. In the case of genotypes, it ranged from the lowest for GN (24.26%) to highest for RDW (63.94%) followed by RL (54.26%). Considering the stress level, it was the lowest for MGT (57.29%) to highest for SL (99.19%) which was closely followed by the rest of the traits. In respect of genotypes and stress kevel interaction, it ranged from 24.35%-92.69%. However, the highest was observed in SFW (92.69%) followed by CEG (91.18%) and VI (90.55%) while lowest was recorded in RL (24.35%) and RDW (26.80). The results exhibited moderate to high heritability (24.26%-99.19%) along with moderate genetic advances for the majority of the characters which offer to chances expected response to selection. The characters which exhibited high heritability indicates that the selection will be more effective whereas the traits showing low heritability indicates that the selection will be affected by the environmental factors [26]. The high heritability of a trait also indicated that this trait was control by the additive genetic effects. This was in line with Sabu et al., 2009 [28] who noticed that the presence of relatively high heritability in a particular trait indicates more additive gene effects in favor of probable improvement. The results of this study revealed that the variability amongst the majority traits was mostly due to genotypic variance though there was a little contribution from the environment on the respective traits. Narayanankutty et al. 2003 [29] studied on thirty-seven vegetable cowpea genotypes for observed the genetic variability and divergence, and they obtained significant differences among the genotypes for all the recorded characters. High heritability coupled with high genetic advance was recorded for seedling dry weight, seedling vigor index of mungbean [22]. Johnson et al., 1955[30] state that heritability coupled with genetic advance would be more dependable and necessary in formulating the selection methods. An estimate of heritability is essential for applying the optimum breeding strategy [26]. Thus, heritability regulate the skill of selection, though the usefulness of selection for a particular trait depends on both the relative extent of genetic and environmental factors which reflex of phenotypic differences among genotypes in a population [26].

### Genetic Advance and Genetic Gain

In the case of genotypes, the genetic advance varied from lowest for MGT (0.09) to highest for RDW (0.41) followed by GN (0.40), SFW (0.37) and VI (0.36), respectively. At the stress level, it ranged from 0.02-0.78. However, the highest was in GN (0.78) followed by VI (0.42) and lowest in MGT (0.02). In respect of genotypes and stress level interaction, it ranged from 0.0-0.42 whereas, maximum (0.42) were recorded both in GN and VI. The lowest (0.1) was in RDW. Similarly, in the case of genotypes, the genetic gain varied from lowest for GN (8.59) to highest for RDW (45.61) followed by SDW (30.88) and RL (30.34). At the stress level, it ranged from 2.20-34.46. However, the highest was in SL (34.46) followed by SG (32.50) and the lowest was in MGT (2.20). In respect of genotypes and stress level interaction, it ranged from 0.34-22.05whereas, maximum (22.05) were recorded in MGT followed by CEG (17.72). The lowest (0.340) was in RDW. Degefa et al., 2014 [26] mention that the minimum value of genetic advance derives from a low score of phenotypic variance or observed as a result of non-additive gene action which have epistatic and/or dominance effects. An increase in the magnitude of genetic advance across the water stress gradient indicated an increase in the inherent variation in the response of genotype to drought and a possibility of selection of suitable genotypes at higher water stress. Comparable observations were also reported earlier study by Allah et al., 2014[31].

### Genetic Diversity Analysis of 33 Mungbean Genotypes

#### The Clustering Pattern and Distribution

Results of variance analysis for individual traits were found significant which was suggesting the need for estimating D2 for further study. As regards, thirty-three (33) mungbean genotypes were assembled into five different clusters using Mahalanobis’s [32] D2 statistic based on their genetic variability present in the respective genotypes (Table 3). Cluster pattern revealed that cluster III was the largest group consisted maximum genotypes (11) closely followed by cluster IV with ten genotypes. Clusters II and V consisted of six and five genotypes, respectively whereas, clusters I contained only one genotype. The genotypes which were accumulated in the same cluster indicating that they were not sharply diversified.

**Table 3.** Distribution of 33 genotypes of mungbean in five clusters

| Cluster | Number of Genotype | Genotypes |
| --- | --- | --- |
| I | 1 | BARI Mung-1 |
| II | 6 | BARI Mung-3, BINA Mung-8, BU Mung-4, BMX-05001, BMX-09009-6 and VC-2764B |
| III | 11 | BARI Mung-4, BARI Mung-5, BARI Mung-6, BARI Mung-7, BINA Mung-6, BU Mung-2, BMX-030011-6, BMX-04005-3, BMX-05009, BMX-050012-6 and BMX-06001 |
| IV | 10 | BINA Mung-5, BMX-01019-1, BMX-01007, BMX-05004, BMX-06007, BMX-07007-4, BMX-07009-10, BMX-08011-8, BMX-08011-2 and BMX-97024-13 |
| V | 5 | BARI Mung-2, BARI Mung-8, BMX-01015, BMX-08009-7 and BMX-08010-2 |

#### Contribution of Different Traits Towards the Divergence of Genotypes (Principal Component Analysis)

Eigen values represented the weight of different traits to the total divergence. it also helps in the assessment of diversity and also was used to expose the relationships more clearly. The contributions of 11 quantitative traits to genetic divergence among the 33 mungbean genotype were assessed by a rank average of the individual trait. The trait contributing to maximum divergence requires more consideration for selection in the hybridization program reported by Varma [20]. The first five components were found to contribute to 98.19 % of the total variation while only the first two components contributed to 82.12 % variation (Table 4). Hence, scores gained from the first two components were plotted against two main axes and after that put on with clustering (Fig.1). Earlier results of D2 analysis confirmed with the pattern of clustering in the plot.

**Table 4.**
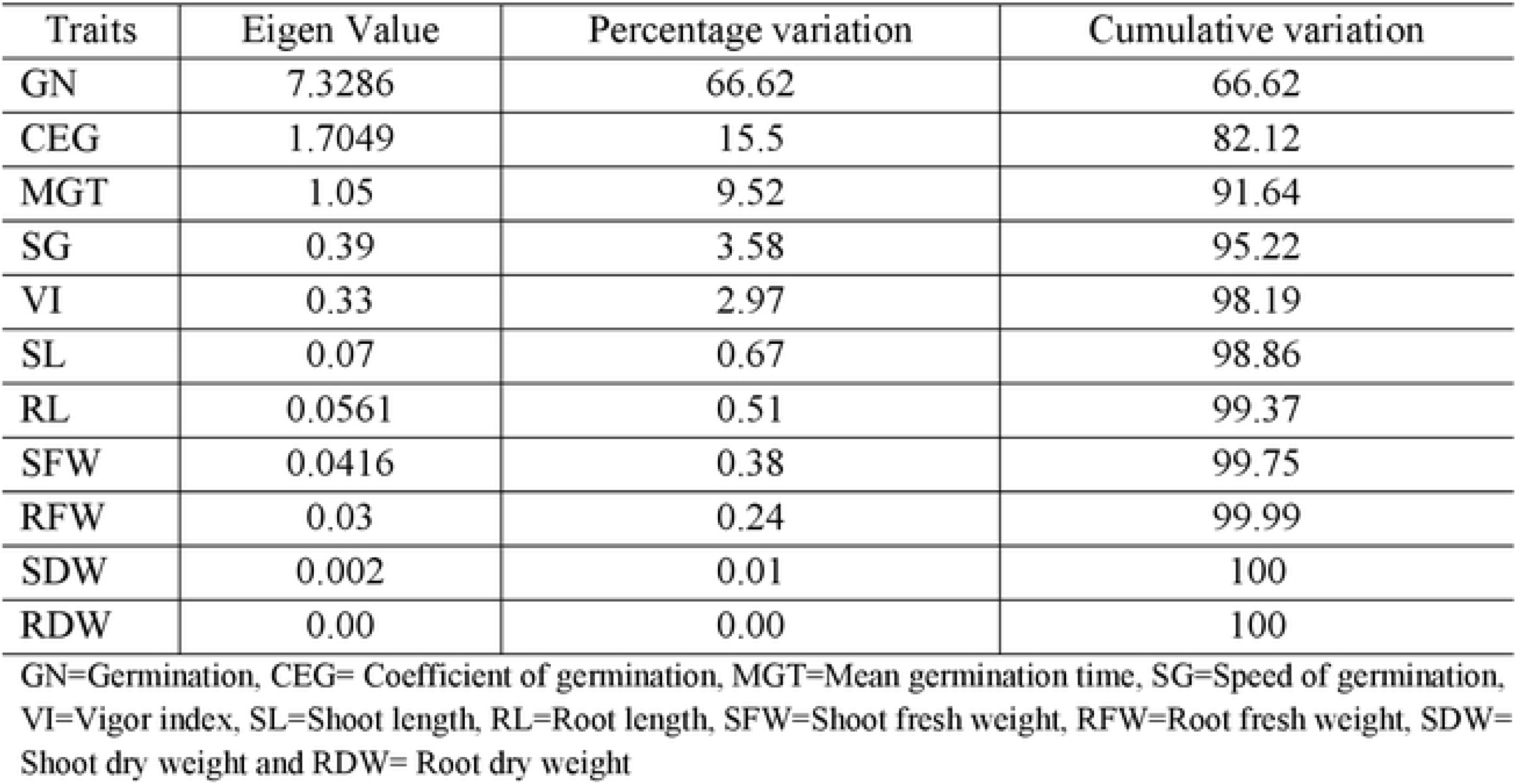
Eigen values and percentage variations contributed by eleven different component traits in 33 mungbean genotypes

**Fig. 1.**
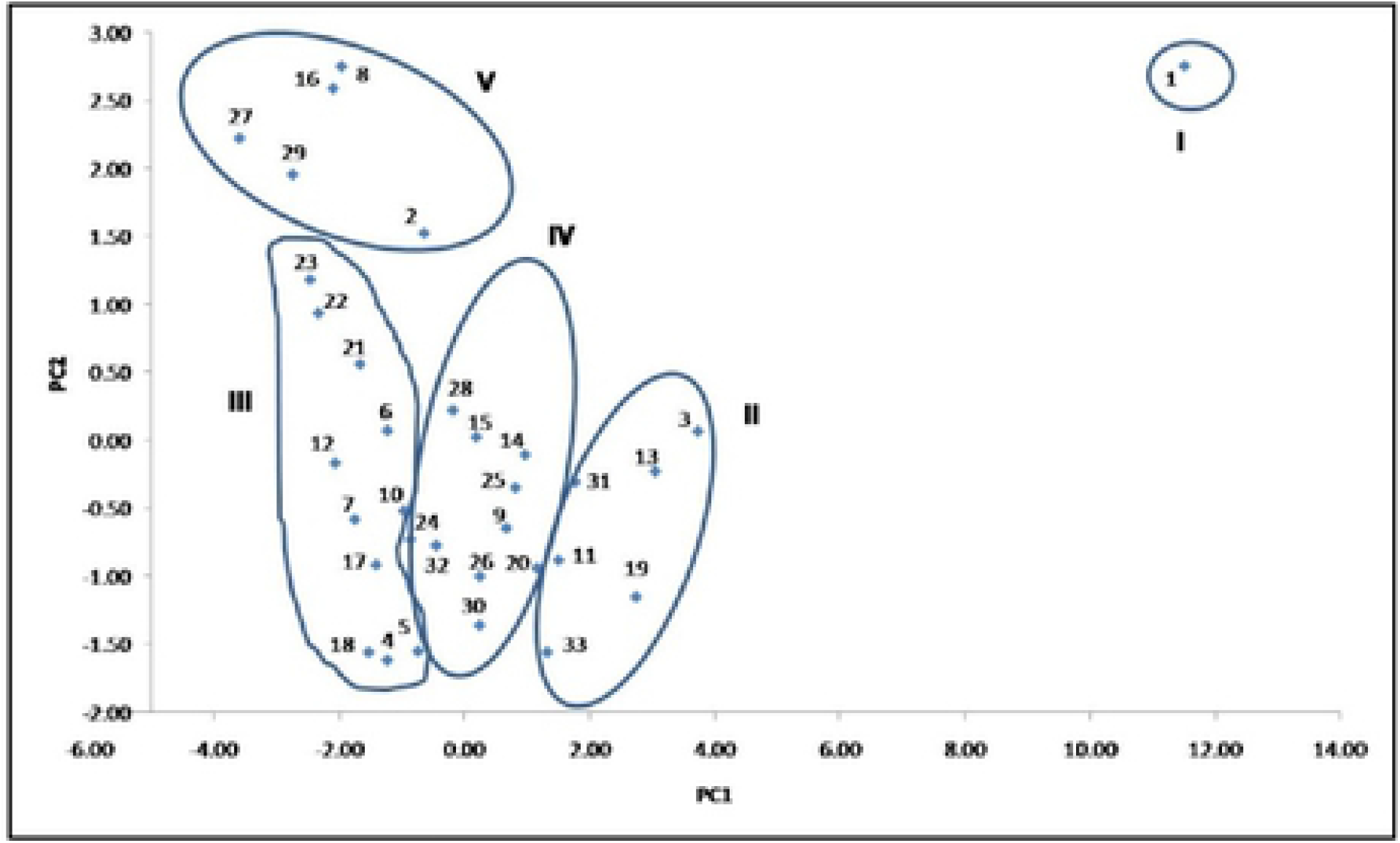
Allocation (scatter)of 33 mungbean genotypes on the basis of their principal component scores superimposed with clustering (1….33=serial no of genotypes)

#### Intra and Inter Cluster Distances (Canonical Variate Analysis)

The intra and inter-cluster distances were estimated to find the distances within and between the clusters containing different genotypes. The average intra and inter-cluster distance values among the five clusters were presented in Table 5. The extent of intra cluster distances indicated the diversity present within the genotypes in the cluster [33]. The inter cluster distance found more than the inter clusters distances explaining that diversity among the genotypes between the clusters was more. It was revealed that the most divergent cluster was cluster I as it was having the highest inter cluster distance compared to other clusters which ranged from 16.23 to 16.5 indicating that the cluster I was distinctly diverse from others and the genotype present in the cluster. However, the highest distance was attained between cluster I & III (16.50) signifying the extensive genetic divergence between these two clusters which was followed by cluster I & II (16.35) as well as cluster I & IV (16.33). The minimum distance was observed between cluster II and IV (2.50) followed by clusters III and IV (2.81). This indicating that genotypes present in those clusters are closely related [22]. Thus, hybridization between genotypes of these clusters may not yield a high level of heterotic expression in the F1’s and variability in segregating progeny (F2) [33]. Hence, Parents could be selected based on larger inter-cluster distance to obtain novel recombinants in subsequent generations [33]. The maximum within-cluster distance (D=0.12) was recorded in Cluster II which was closely followed by cluster V (0.11) and cluster IV (0.10), and which indicate the existence of maximum differences among the genotypes that fall in these clusters [22].

**Table 5.** Average Intra and inter cluster distance (D2) of mungbean genotypes

| Cluster | I | II | III | IV | V |
| --- | --- | --- | --- | --- | --- |
| I | **0.00** |  |  |  |  |
| II | 16.35 | **0.12** |  |  |  |
| III | 16.50 | 5.01 | **0.09** |  |  |
| IV | 16.33 | 2.50 | 2.81 | **0.10** |  |
| V | 16.23 | 8.04 | 4.22 | 5.56 | **0.11** |

The cluster mean values of the eleven different traits were given in Table 6. It was observed that cluster I gave the lowest mean values for all the traits except mean germination time. Cluster III had the maximum mean value for SFW, SDW, RFW and RDW. However, the genotypes in the cluster V recorded for greater performance by giving the highest mean value for germination, coefficient of germination, speed of germination, vigor index, shoot length and root length. This comparison indicates that cluster III and V had better cluster means for most of the characters. Hence, cluster III and V might be considered better for choosing the parental genotypes used for hybridizing with the counterpart from cluster I those may generate novel recombinants with anticipated characters. These is also agreement with the findings of Garg, et al., 2017; Shweta, 2013; Uz-Zaman et al., 2013; Singh et al., 2009; Sarkar and Kundagrami, 2016 [22, 34, 35, 36, 37].

**Table 6.** Clusters means of different characters in mungbean genotypes

| Cluster | GN (%) | CEG (%) | MGT (days) | SG (No./day) | VI | SL (cm) | RL (cm) | SFW (mg) | RFW (mg) | SDW (mg) | RDW (mg) |
| --- | --- | --- | --- | --- | --- | --- | --- | --- | --- | --- | --- |
| I | 52.00 | 14.90 | 5.44* | 2.04 | 874.07 | 6.76 | 5.43 | 63.80 | 23.19 | 6.84 | 3.59 |
| II | 62.72 | 18.68 | 5.41 | 2.48 | 896.95 | 6.37 | 6.19 | 70.74 | 39.97 | 8.00 | 6.87 |
| III | 68.64 | 19.44 | 5.19 | 2.80 | 1275.12 | 8.90 | 8.11 | 123.33 | 56.66 | 13.13 | 10.20 |
| IV | 69.77 | 19.09 | 5.30 | 2.79 | 1000.07 | 6.67 | 6.53 | 96.80 | 44.66 | 10.21 | 8.01 |
| V | 75.60 | 20.99 | 4.81 | 3.31 | 1554.95 | 9.76 | 9.84 | 98.90 | 45.76 | 10.43 | 6.67 |

GN=Germination, CEG= Coefficient of germination, MGT=Mean germination time, SG=Speed of germination, VI=Vigor index, SL=Shoot length, RL=Root length, SFW=Shoot fresh weight, RFW=Root fresh weight, SDW= Shoot dry weight and RDW= Root dry weight, \*Average of four stress level data as the genotype under cluster I failed to germinate at -4 bar stress level

To sum up, a number of 33 mungbean genotypes were imposed with drought stress induced by PEG solution. Variability and diversity were studied as influenced by drought stress treatments. Regarding the results significantly higher PCV compare to GCV was found for all the traits, representing that these traits was influence by the environmental factors. In addition, the heritability was showed moderate to high (24.26%-99.19%) among the study traits indicating these traits were controlled by additive genetic effects. The clustering pattern exhibited that cluster III were contained maximum eleven genotypes followed by cluster IV with ten genotypes. Cluster I, II and V consist of one, six and five genotypes, respectively. In the case of a contribution of different traits, the first five components were found to contribute to 98.19% of the total variation while only the first two components contributed to 82.12% variation. The maximum inter-cluster distance was recorded between cluster I with rest of the clusters. The least was obtained between cluster II and IV. On the contrary, the highest intra-cluster distance was observed in cluster II (0.12) followed by cluster V (0.11). The genotypes in the cluster V recorded for greater performance by giving the highest mean value for germination, coefficient of germination, speed of germination, vigor index, shoot length and root length. This means that genotype under cluster V has any gene or mechanism which responsible for a tolerant higher level of water stress. Selection of parental genotypes from these clusters might resulted higher heterosis if hybridized. The outcome of the study could be helpful for selection water stress-tolerant mungbean genotypes under breeding program.

